# Calming Meditation Increases Altruism, Decreases Parochialism

**DOI:** 10.1101/060616

**Authors:** Karl Frost

## Abstract

It has been proposed that cultivating calm will increase altruism and decrease parochialism, where altruism is defined as self-sacrifice in support of others, regardless of group affiliation or identity, and parochialism is defined as prosocial self-sacrifice restricted to fellow members of a group. Such could be the case with a calming meditation practice. An alternate hypothesis, coming from the study of ritual, proposes that shared practices lead to bonding, increasing parochialism, but not altruism generally. These contradictory hypotheses of the potential effects of shared cultural practices of calming meditation were explored via a formal behavioral experiment using a simple treatment and control format with a short, facilitated breath awareness practice known to produce calm. Altruism and parochialism were measured through anonymous play in Public Goods games performed with both in-group and out-group individuals. The sum of contributions of the two plays gave a measure of altruism, while the difference between the two gave a measure of parochialism. The analysis of the results using Bayesian AICc model comparison methods supports the first hypothesis that calming practices reduces parochialism and increases altruism. The hypothesis of intentional shared practice as parochialism inducing was not supported by the results in this case of a shared calming practice.

## Introduction

The major religions of the world all hold prosociality as sacred, contextually promoting altruism, compassion, generosity, honesty, trust, cooperation, forgiveness, etc.˙ Examples abound from Buddhism and Hinduism to teachings in the Abrahamic religions (Judaism, Christianity, Islam). What it means to be part of a religion is conveyed partially through text, but also through physical practices and rituals. Questions arise as to whether practices within these religions cultivate prosociality independent of the text and whether this prosociality is directed evenly to all or limited to fellow group members. For the purposes of this paper, ‘generalized altruism’ will refer to willingness to practice self-sacrificing behavior that benefits others, where the others may or may not be members of the same group, while ‘parochialism’ will refer to the willingness to practice such altruistic behavior, but only with members of some in-group. These are not mutually exclusive; an individual might be somewhat altruistic, but also be parochial, willing to be generous toward outsiders, but more generous toward fellow group members. Calm, contemplative practices within these religions and in secular contexts are often motivated in part by the cultivation of universal compassion: Buddhist loving kindness meditation (LKM), the instructions for ritual in association with compassion throughout the Quran, the ritual of Sunday sermon in Catholicism, etc.. This is then an empirical question: do calming practices lead to general altruism, and if so, is this effect independent of explicit prosocial rhetoric? Similarly, do such practices reduce parochialism, and if so, is this effect independent of prosocial rhetoric?

A counter hypothesis that is often posited in the anthropological literature is that rituals, particularly when performed as a group, exist because they functionally bond groups together into communities of shared norms and cooperation (Sosis & Alcorta, 2003). This would allow groups who share these practices to be able to overcome public goods type cooperation problems and thus thrive. By this hypothesis, such group ritual activities will increase parochialism, rather than general altruism. It is a standing question whether behaviorally a shared practice of calming meditation would be best described as 1) a general altruism enhancer, 2) a parochialism enhancer, or 3) a neutral practice with no effects on altruistic behaviors.

Altruism is a long standing problem in evolutionary theory (Fehr & Fischbacher, 2003). While altruism provides a public good it is, by definition, at the expense of a private good. This is characterized by the social dilemmas of the Prisoner’s Dilemma or Public Goods. While groups of individuals who sacrifice of themselves for the greater good would transparently do better than groups of selfish individuals, such groups are vulnerable to free-riders who take advantage of the public goods provided by others’ sacrifices without themselves sacrificing. There are group level incentives to act one way, but individual level incentives to act in a contrary way. Lacking some sort of policing or other mechanism for the maintenance of adherence to altruistic norms, such selfish individuals do better than altruists within the cooperative group and evolutionary processes undermine altruism through the success of such selfish individuals.

There are many proposed theoretical mechanisms for the maintenance of altruism in groups of unrelated others which have been shown to contextually stabilize or promote altruism. Repeated play and reciprocal altruism can do this (Axelrod & Hamilton, 1981; Trivers, 1971). Policing and the ability to punish non-cooperators has been shown to stabilize altruistic norms (Henrich & Boyd, 2001). Still, there are many situations in which behavior is not under reliable surveillance and therefore neither reciprocity nor policing can stabilize altruism. Despite this, humans have been shown cross culturally to be able to maintain altruism in such anonymous contexts (Henrich et al., 2005). If shared practices are able to induce bonding, they can in some circumstances evolutionarily stabilize altruism (Frost, 2016). If, however, a shared practice (as a calming meditation) induces more general altruism, this represents an evolutionary dilemma; where there is some other factor causing preferential assortment of such altruists (such as preference for interaction with fellow ritual practitioners) such altruism may be promoted through cultural group selection, (Wilson, 2002). Perhaps this may be the case with such calming practices, if they do promote more generalized altruism. Alternatively, the solution to the evolutionary puzzle could involve associated independent benefits of the practice, countering any losses to freeriding in public goods type situations. In any case, this is an issue of the cultural evolutionary trajectory of such practices and should be seen as an independent question from the observable effects of the practice.

There is a great public interest in meditation practices as well as a growing body of scientific research on the physiological and psychological impacts of meditation. This has included investigation into the prosocial implications of meditation and related calming practices, like yoga, Tai chi, and prayer, suggesting that calming ritual practices promote altruism. Meditation training was found to result in increased helping behaviors toward strangers in need (Condon, Desbordes, Miller, & DeSteno, 2013). Brief loving kindness meditation (LKM) and related compassion training was found to generate significantly increased levels of prosocial sentiment (Hutcherson, Seppala, & Gross, 2008; Weng et al., 2013) as well as more altruistic behavior as measured through offers in economic games (Leiberg, Klimecki, & Singer, 2011; Reb, Junjie, & Narayanan, 2010). Yogic meditation was likewise found to increase trust in economic games (Bartolomeo, Papa, & Bellomo, 2012).

Of course, not all meditation practices are the same. Offering prosocial sentiment is different from mindfulness cultivation is different from attention focusing, which is again different from a simple relaxation. It might be expected that contemplation of prosocial sentiments might have effects on prosociality independent from other practices. In a comparing mindfulness practices to simple relaxation practices, indistinguishable effects were found on altruism as measured by play in economic games, indicating that perhaps the effects of mindfulness on altruism are mediated largely through relaxation (Tappen, 2013). A number of metastudies of the physical and psychological benefits of meditation, found common effects amongst a variety of meditation practices mutually characterized by comfortable body position or movement, focus of attention, open attitude: they lead to calm and stress reduction, implying that many of their effects may be mediated through calm (Arias, Steinberg, Banga, & Trestman, 2006; Greeson, 2009; Holzel et al., 2011; Horowitz, 2010). Calming video games have similarly been shown to be correlated with increases in prosociality across multiple measures, while exciting video games have shown the opposite effect (Whitaker & Bushman, 2011). While confounds still exist, these studies together suggest that calm-inducing practices may increase altruism.

Research on attachment and meditation suggest that meditation leads to secure attachment (Sahdra et al., 2011). Attachment security, whether by disposition or induced by treatment, has been shown to predict higher levels of altruism and reduced parochialism (Mikulincer & Shaver, 2005)(Mikulincer & Shaver, 2007)(Mikulincer & Shaver, 2001). As attachment security is a measure of calm in the face of relationship stress, these studies further support the idea that calm increases prosociality and reduces parochialism.

Studies which look at increases in cooperation solely with anonymous others did not however compare in vs out group cooperation levels, and so do not independently test parochialism. One study attempted to assess impacts on parochialism and found that daily spiritual experiences (meditation, prayer, mindfulness, and spiritual experiences) were associated with higher levels of volunteering, charitable donations and helping behavior, but that they better predict helping unrelated others than helping friends (Einolf, 2011). This study did not single out ‘calm’, however, and there are many potential confounds. Also, both a brief guided mindfulness meditation (Lueke & Gibson, 2015) and low intensity 6 week compassion meditation training (Kang, Gray, & Dovidio, 2014) were found to decrease negative out-group racial biases. This supports a more general hypothesis of reduction in parochialism for these different meditation practices, at least in terms of sentiment, if not behavior.

Mostly disconnected from the empirical study of meditation practices has been a growing scientific interest in the evolution of religion and ritual practices related to group bonding (Atran & Norenzayan, 2005). Foci of investigation include the genetic behavioral predispositions engaged by religion and ritual, cultural evolution of variants in religious practice, and implications of evolutionary dynamics of religion on social structure (Guthrie, 1993; Kirschner & Tomasello, 2010; Shariff, Norenzayan, & Henrich, 2010; Whitehouse, 2004). Durkheim’s definition of ritual is the canonical one in anthropology and sociology: behaviors whose economic or survival function is opaque (Durkheim, 1912). However, this category has more to do with the observing anthropologist and what they subjectively consider opaque than with functions or objective qualities of the observed behaviors. As such, ‘ritual’ remains a problematic term, a kind of ‘grab bag’ of different behaviors, and not necessarily a clear category in terms of intrinsic qualities and function of the behavior in cultural context. For the purposes of this investigation, I look more narrowly at those specific shared practices where anthropologists propose a group bonding function. These include shared sacrifice (Ruffle & Sosis, 2007), intentionally visually synchronized movement (McNeill, 1995), and shared behavioral norms (Whitehouse, 2000). As a shared practice of calming meditation involves both a shared sacrifice of time and intentionally synchronized or shared norms of practice, it is a question, whether a shared calming practice cultivates parochialism and not altruism.

Central to anthropological theories of the cultural evolution of some forms of ritual is their ability to help groups public goods type dilemmas and free rider problems (Sosis & Alcorta, 2003; Wilson, 2002). Academics, starting with Durkheim and through Rappaport and into the current wave of quantitatively-minded scholars, explore how religion and ritual can be group level adaptations that coordinate individuals for more effective collective action within groups and how socially learned systems of prosociality spread through cultural group selection (Atran & Henrich, 2010; Durkheim, 1912; Norenzayan & Shariff, 2008; Rappaport, 1999; Richerson et al., 2014; Wilson, 2002). A theme of many anthropological studies of ritual is that rituals are bonding practices that facilitate parochialism. The costly signaling theory of ritual practice suggests that shared sacrifice acts as a marker of commitment to group norms (Irons, 2001). It suggests that rituals only benefit in-group, not out-group prosociality, and only when done as a signal with witnesses, as opposed to private rituals. This has been supported through observed correlations between longevity and costly ritual displays in utopian communes(Sosis & Bressler, 2003) and observed increases in parochial behavior in economic games in religious vs secular kibbutzes (Sosis & Ruffle, 2003). Suggesting that this effect may be instinctual and independent of intention, simply sharing stressful experiences has been found to lead to group prosociality in a study of economic games amongst people living in evacuation shelters post Hurricane Katrina in Houston, Texas (Whitt & Wilson, 2013). Alternatively, McNeil hypothesized that we instinctively have prosocial sentiments toward those with whom we have successfully synchronized or coordinated in some activity. Synchronizing and coordinating movement with others, as in unison dance or coordinated work, has been observed to promote altruistic feelings with task co-participants(Kirschner & Tomasello, 2010; McNeill, 1995; Rappaport, 1999), though this effect is most pronounced when this synchrony is intentional rather than a coincidence or coordinated externally (Reddish, Fischer, & Bulbulia, 2013). These observations support the theory that ritual helps define group boundaries and/or engender prosocial sentiments toward fellow group members, and altruism will increase toward these fellow group members, but not toward out-group individuals. Studies do not all agree however, as at least one study has shown that synchrony can lead to more generalized rather than parochial prosociality (Reddish, Bulbulia, & Fischer, 2013). This indicates that this is an open question and that perhaps parochial effects of synchrony may involve synergy with other factors, like ‘dysphoric’ sacrifice (Whitehouse, 2000). Whitehouse and colleagues have observed that ritual practice as documented in ethnography seems to fall primarily in one of two modes: *imagistic* mode rituals which are highly arousing and infrequently performed (like endurance or pain oriented ceremonies) and *doctrinal* mode rituals which are more frequently performed and are less arousing (like daily group prayer). Such imagistic rituals are theorized to create strong bonds in small groups where doctrinal mode rituals create lighter bonds in much larger groups, which matches observed correlation between size of society and ritual practice in the ethnographic record (Atkinson & Whitehouse, 2011). This suggests an increase in scope of prosocial relations as the arousal level of the ritual decreases.

Given the heterogeneity of behaviors that are called ‘rituals’, instead of a unified theory of rituals, it is more plausible that we will find separate theories of the effects of synchronous movement, calming, dysphoric practices, ecstatic practices, repetition, attention focusing etc., perhaps with synergistic effects. Where it might be that something like a stressful synchronous dance activates a native behavioral disposition toward bonding and parochialism, it is perhaps also the case that we have instincts to be altruistic when we are in a relaxed and unstressed state. This could have evolved, for example, if an inner state of calm was consistently correlated with being in a state of lower resource stress and with being around relatives, where we would have had low cost for helping kin. Artificially inducing calm through ritual practices could hijack this evolved psychological disposition to cooperate, activating it in novel contexts. Whether or not this is an accurate depiction of the evolutionary origin of an instinctual response of altruism and reduced parochialism in humans, a growing body of research, including this study, explores this hypothesized effect.

This study attempts to answer the question of whether calming practices are best modeled as promoters of general altruism, parochialism, or neutral. It does so via a formal behavioral experiment using a simple treatment and control format with a short calming breath awareness practice, with altruism and parochialism measured through anonymous play in public goods games performed with both in-group and out-group individuals. This study aims to further our understanding by assessing the effects of calming meditation simultaneously on in-group and out-group prosociality. This allows for a comparison of bonding theories and the calming theory to see which better predicts resulting changes in altruism and parochialism. Three hypotheses were compared in this study to see which hypothesis best predicts the behavioral effects of a socially learned calming practice. The first hypothesis (Calming Hypothesis) is that calming meditation leads to an increase in altruism and a decrease in extant parochialism. The second hypothesis (Group Bonding Hypothesis) is that calming meditation, as a bonding ritual, will lead only to an increase in parochialism, and only when performed as a group. The third hypothesis (Null) is that calming meditation has no effect on altruism or parochialism.

## Method

### Participants

The experiment was performed at the beginning of each night of an interactive theater work at UC Davis over 7 evenings in February 2011 (Frost, 2011). The fact that the performance was beginning with a formal behavioral experiment was included in publicity materials for the show, but the advertised theme of the performance did not give any cues about the content of the experiment. While one might expect those who attend a theater piece to be more generally open to new experience, the comparison between treatments and random assignment would be expected to eliminate any self-selection issues due to it being in a theater piece. The results of those who had already participated in previous evenings of the performance, who had previously studied the psychology of Public Good games, or who did not want to play the Public Goods games were excluded.

The included participants (n=331) were a mixture of Davis and Sacramento theater goers and UC Davis students, with treatment groups ranging from 10 to 16 in size. Ages ranged from 17 to 65, and overall 149 male and 182 female. As participants entered the theater space, they were given a card at random with a letter and number on it, which was used to indicate their treatment group and to identify them for any winnings at the end. They were then asked to follow facilitators into different spaces for treatment and testing. Winners were announced at the end of the evening. After each performance audience members were invited to share and discuss their experiences of decision-making during the experiment, as well as discuss the broader social and political issues of cooperation and altruism. This post-performance feedback was used to contextualize and interpret the results and to informally assess understanding of the game structure. Part of the motivation for this study group was to expand beyond the typical study population of college students.

### Procedure

The experiment followed a randomized treatment and control group design to test for the effects of a calming meditation on altruism and parochialism. Participants were randomly divided into one of four equal sized groups: two control and two treatment groups. After treatments all groups anonymously played two rounds of a Public Goods game (Dawes, 1980), once with members of the treatment group and once with a random mixture of members of all groups. The total offers in the both plays of the game were used as a measure of altruism. The difference between in-group and out-group offers was used as a measure of parochialism.

Participants in one treatment group (*Solo Meditation*) were brought together into a room, asked to face a blank wall away from each other, and facilitated in a 5 minute breath awareness meditation with their eyes closed before play. The second treatment group (*Group Meditation*) was facilitated in the same meditation, but oriented in a circle facing each other, with participants asked to look at each other before closing their eyes for the meditation. The breath awareness practice was adapted from preparatory theater training exercises known to regularly produce calm and was similar to that in Mindful Breathing treatments in other studies (Arch & Craske, 2006; Feldman, 2011), but with less emphasis on attention refocusing and with instructions for allowing a relaxed stance and breath. This deemphasized the potential confound of attention focusing and more strongly emphasized relaxation. See the Supplementary Material for scripts.

The purpose of the Group Meditation treatment was to establish intentional visual synchronization as a group, reflecting the assumptions of the synchrony model of ritual (Hypothesis 2) which predicts that such intentional visual synchrony leads to parochialism amongst practice co-participants. One control group (*No Treatment*) played the economic games immediately. The other control group (*Socialize*) was invited to wait and socialize as they like for 5 minutes before the public goods games. *Socialize* was added in order to control for effects of simply spending time with each other vs being asked to engage in an intentional activity. Facilitators followed scripts and were rotated between roles in each run of the experiment, so that their individual personality differences would be less likely to influence the results. The two meditation scripts were identical except for the instructions to face each other and look at each other first vs instructions to face away from the group toward a wall (to reduce feelings of visually synchronizing with each other).

In an effort to minimize the effects of preconceptions about such concepts as “group”, “cooperation”, “trust”, or “meditation”, the scripts did not reference any of these or related terms. The instructions for the breath awareness meditation were simply for participants to close their eyes, relax, and be aware of their breath, as opposed to asking people to “meditate” or to cultivate any other social feeling, like ‘compassion’ or ‘mindfulness’. Ideas of kindness, compassion, and mindfulness are often layered into culturally situated meditation practices, like Loving Kindness Meditation. We expect that there would be synergistic effects of such embedded messages, but the purpose of the experiment was to isolate the effects of a simple calming practice from any effects of focusing the attention specifically on altruistic concepts.

### Measures

The game played was a version of Public Goods. Public Goods (PG) is the multi-person version of the classic Prisoner’s Dilemma problem. Such games are characterized by the best option for the group being for everyone to cooperate, where individual level incentives motivate people to defect (not cooperate) (Ledyard, 1995). In the classic example, two criminals are arrested together. If neither snitches on the other, they both will get X years in jail. If one snitches and the other does not, the snitch gets out free, but the non-snitch gets Z years. If both snitch, they both get Y years. 0<X<Y<Z. There is a self-centered motivation to snitch, because whether the other snitches or not, it is always better for oneself to snitch. The altruistic option is to cooperate. It is altruistic in that the group does better with cooperation, but the individual would do better with defection. Such coordination problems cause groups of selfish individuals to all defect, leading to bad outcomes for all individuals, and thus the group. Public Goods has the same payoff structure, but involves more than two people. It has the additional problem of increased potential anonymity, when one doesn’t know who played how.

In this version of Public Goods, each participant was given 10 points. They could contribute as many points of the 10 as they wanted to a pool and keep the rest. In each round, one player would win a sum of money. The amount won was proportional to the number of points put into the pool up to $20 if all points were put in. Each player’s chance of winning was proportional to the number of points they kept, plus one. Thus, a player was most likely to win if they kept all of their points, but the pot would be biggest for whomever got it if everyone put in all points. In the first round of the public goods game, participants played with members of their treatment group, while in the second round, they were asked to play with a similar sized group randomly drawn from all groups (and thus on average having 75% participants from other groups). All plays in both rounds were anonymous. The number of points contributed altogether was used as a measure of altruistic trust, one of the dependent variables. The difference between in-group offers and out-group offers was used as a measure of parochialism, the other dependent variable. While groups were randomly formed and there were no explicit cues given about cooperation, competition, or group identity, simply separating people randomly into groups has been shown to reliably induce functional group identification and parochialism (Dawes, 1980) (Kollock, 1998). If the control group individuals demonstrated parochial play in the PG games, this would demonstrate that the experimental set up successfully induced groups behaviorally relevant for altruism decisions. A change in the in-group vs out-group difference in play for treatment groups would indicate a change in parochialism.

An option to ‘Not Play’ was added. It was found in test runs that a few audience members (humanities graduate students) objected to the idea of the experiment, yet felt they wanted to go through the it and gave nonsense answers. We found that they would refrain from nonsense answers if given the option to answer ‘No play’.

Winners were not announced until later in the evening, to minimize ordering effects in play in establishing parochialism. Further, if there was an ordering effect, this would not change findings, since the order was constant between treatments and the difference between groups with regards parochialism is relevant, not the base level. Subjects were asked to not discuss the experimental set up with those whom they knew were attending on subsequent evenings. A brief demographic survey was added after the game play. All answers from those who replied that they had studied economic games or had participated in the performance on another evening or who answered ‘No Play’ in the games were discarded.

See Supplementary Material for experimental scripts.

### Data Analyses

Bayesian AICc model comparison methods were used to analyze the data, using ordered logit regression models to assess the predictive relevance of including various independent variables (Burnham & Anderson, 2002). Bayesian methods were chosen over the currently more commonly used Fisherian methods of p-value testing. This choice of data analysis was used because of the better fit of Bayesian methods to the analytic task of assessing the predictive ability of hypotheses (Gigerenzer, Krauss, & Vitouch, 2004). The p-value is a “measure of the probability of the data, given the hypothesis” and is NOT a “measure of the probability of the hypothesis, given the data”, which is the principled measure to use in assessing the relative usefulness of hypotheses in predicting future observation, a fact that has been pointed out by statisticians critical of standard practice use of p-values for decades. This is an unfortunate misunderstanding that has been shown to be prevalent globally amongst professional scientists (Badenes-Ribera, Frias-Navarro, Iotti, Bonilla-Campos, & Longobardi, 2016; Haller & Krauss, 2002; Lecoutre, Poitevineau, & Lecoutre, 2003). The misunderstanding and misapplication of the p-value is why some journals are starting to ban its use (Siegfried, 2015). Also, while the p-value is not a principled way to compare two hypotheses (a null and proposed hypothesis) as it is currently normatively used, it is even less useful in simultaneously comparing several hypotheses, which model comparison methods allow and which is required for this study (McElreath, 2012). As has long been pointed out, such simultaneous comparison of all plausible available models is necessary in order to avoid biasing toward favored theories (Chamberlin, 1897).

To interpret AICc results, a model weighting of X% through AICc implies a X% chance that a model (theory) would perform the best amongst compared models at predicting outcomes, given the priors. In other words, if hypothesis 1 is weighted at 10%, hypothesis 2 is weighted at 20%, and hypothesis 3 is weighted at 70%, then the observed data imply this chance (10%/20%/70%) that the model implemented from the hypothesis will be the best amongst the 3 tested hypotheses in predicting future observations, dependent on prior assumptions (taken to be even in this case, ie no biasing). 95% confidence intervals are provided for estimated model parameters to have a sense of the range of most likely effect sizes.

For the data analysis here, first all treatment groups were analyzed together to explore if there was a reliable effect of group formation and parochialism based on division into treatment groups, by seeing if in vs out-group play was heavily weighted by AICc comparison. If this was the case, then the experimental design would have successfully triggered parochial group identification. Next, both total offers and difference between out-group and in-group offers were analyzed as dependent variables to see if AICc weighted significantly the *Meditation* treatments and/or the *Socializing* control in models for either dependent variable.

Bonding theories predict only an effect for the group meditation, which would be to increase both total offers (altruism generally) and the difference between in-group and out-group offers (parochialism). These theories would predict effects for neither the solo meditation nor the socializing treatment. The calming theory predicts that prosocial benefits would result from both solo and group meditation, increasing total contributions and decreasing rather than increasing the difference between in-group and out-group offers. In terms of the AICc model comparison, heavier weighting of models with regression terms for group meditation and with positive regression parameters for both measures would support the bonding hypotheses. Heavier weighting of models with regression terms for both meditation treatments, but with negative intercept for the parochialism models and positive intercepts for the general altruism models would support the calming hypothesis. Dominant weighting for the null model would, of course, support the null.

## Results

Figures 1A–1C give the observed mean offers for public goods game play for all 4 treatment groups: in-group, out-group, total offers (as a measure of general altruism), and the difference between in and out group (as a measure of parochialism). While the effects were small, as might be expected of a short, one-off, treatment, the results better support the calming model than synchrony or null models, suggesting that calming practices lead to an increase in altruism generically and decrease parochialism.

**Figure 1:**
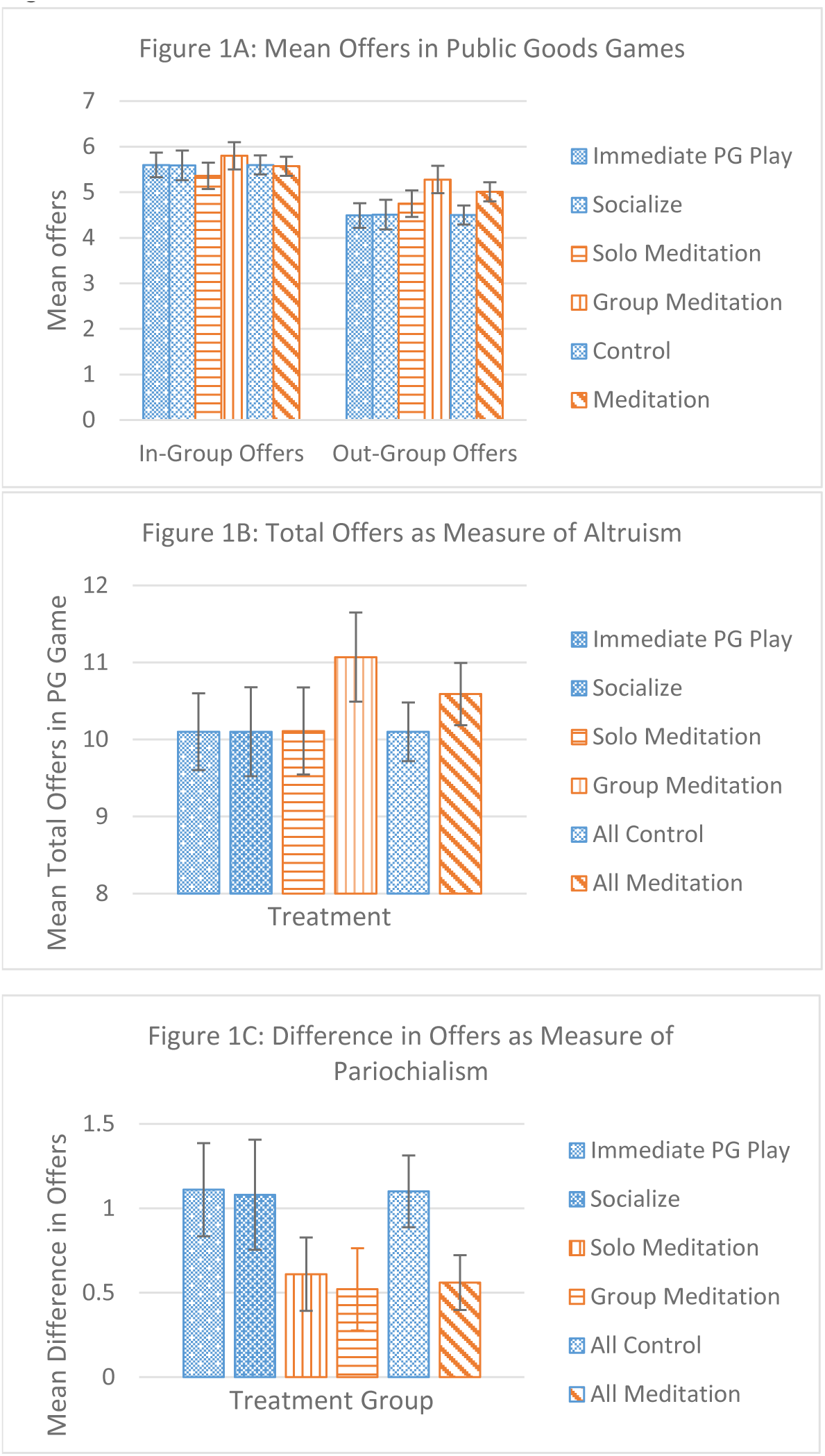
Mean Offers in Public Goods Games *(with standard errors)*

Visual examination alone shows that there was a significant difference between in-group and out-group offers overall and particularly in the controls. The experimental set up clearly triggered identification of the treatment group as a group relevant to altruistic decisions and parochialism as explained in the methods section. AICc model comparison selected the model that included a term for in vs out-group play, with a strength of over 99.9%.

Table 1 shows the results of AICc model comparison for both general altruism and parochialism. As can be seen from the table, the null is clearly less weighted than the models tracking meditation treatments. Models with meditation treatments as an independent variable are weighted collectively at 69% for predicting total offers and 75% for predicting parochialism. Table 2 shows the model intercepts for the model tracking both solo and group meditation for the ordered logit regressions and also translates them into odds ratios for increased contributions or increased difference in contributions between the two games. The results are illustrated with 95% confidence intervals in Figure 2. These parameters indicate that the effects of the group meditation on the public goods game play were better characterized by the calming hypothesis than by the synchrony or null hypotheses. While 0 is still just inside the range of the 95% confidence interval for the intercept, it is only barely so, and most of the confidence interval is characterized by positive effects on total offers and negative effects on parochialism. Specifically, looking at the parameters for group meditation, 0 is excluded by the 92% confidence interval for the model of parochialism and the 86% confidence interval for the model of general altruism.

**Figure 2:**
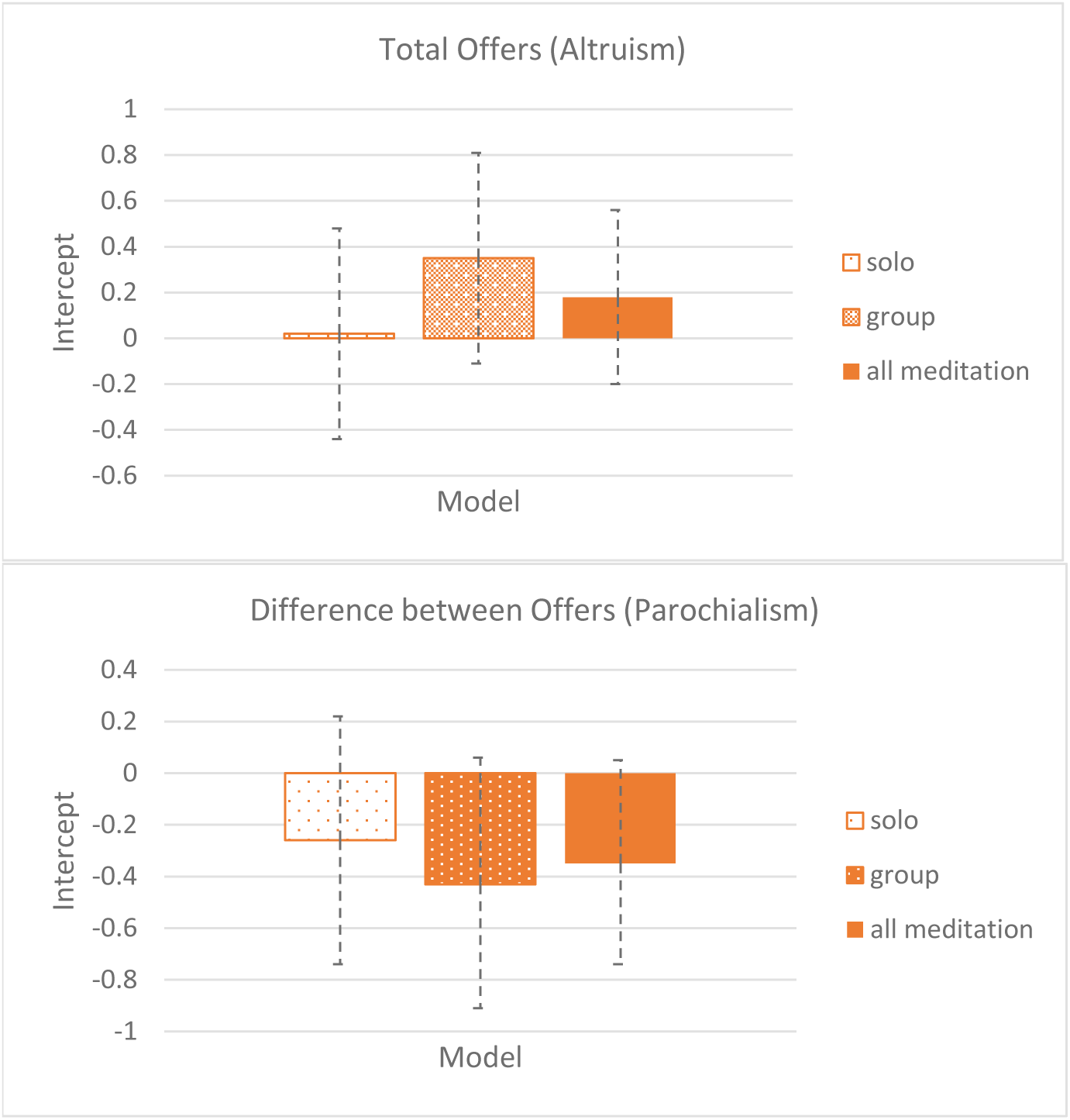
Intercepts for Models with linear relationship between meditation treatment and total offers and offer differences *(plus 95% confidence intervals)*

**Table 1:** AICc model comparison table.

| Model | Parameters | Total offers | Parochialism |
| --- | --- | --- | --- |
| 0 | (none) | 0.23 | 0.17 |
| 1 | Socialize | 0.09 | 0.09 |
| 2 | Solo | 0.08 | 0.06 |
| 3 | Group | 0.24 | 0.16 |
| 4 | Socialize and Solo | 0.03 | 0.03 |
| 5 | Solo and Group | 0.08 | 0.09 |
| 6 | Socialize and Group | 0.08 | 0.06 |
| 7 | Socialize, Solo, Group | 0.02 | 0.03 |
| 8 | Meditation | 0.12 | 0.24 |
| 9 | Meditation and Socialize | 0.04 | 0.08 |
| **All Models with Meditation: Total** |  | **0.69** | **0.75** |

This table lists the AICc model weights for 10 different models which have all combinations of treatment groups as independent variables. “Socialize” refers to the treatment group that was invited to ‘hang out’ for 5 minutes before playing economic games. “Solo” refers to the treatment group asked to perform the breath awareness exercise facing away from each other. “Group” refers to the treatment group that performed the breath awareness practice facing each other. “Meditation” refers to membership in either “Solo” or “Group” treatment.

**Table 2:** Intercepts and Odds Ratios.

|  | Intercept Estimate | standard error | 95% confidence |  | Odds Ratio | 95% confidence |  |
| --- | --- | --- | --- | --- | --- | --- | --- |
|  |  |  | 2.50% | 97.5% |  | 2.5% | 97.5% |
| Difference between In and Out Offers (Parochialism) |  |  |  |  |  |  |  |
| solo | -0.26 | 0.24 | -0.74 | 0.22 | 0.77 | 0.47 | 1.24 |
| group | -0.43 | 0.25 | -0.91 | 0.06 | 0.65 | 0.40 | 1.06 |
| meditation | -0.35 | 0.2 | -0.74 | 0.05 | 0.70 | 0.48 | 1.05 |
| Total Offers (Altruism) |  |  |  |  |  |  |  |
| solo | 0.02 | 0.23 | -0.44 | 0.48 | 1.02 | 0.64 | 1.62 |
| group | 0.35 | 0.24 | -0.11 | 0.81 | 1.42 | 0.90 | 2.25 |
| meditation | 0.18 | 0.19 | -0.19 | 0.56 | 1.20 | 0.83 | 1.75 |

This table lists the intercepts for the ordered logit regression for the meditation treatment terms for models 7 (group and solo tracked separately) and 9 (all meditation treatments combined).
The ‘Odds Ratio’ is the odds of an offer (or offer difference level) or higher, given the treatment divided by the odds of an offer level or (offer difference level) or higher in absence of treatment.

The solo meditation results are harder to interpret. The intercept for total offers is nearly zero, but the intercept for parochialism is, as with group meditation, negative. While the lack of a net effect on total offers would seem to support the null hypothesis, the negative intercept for parochialism supports the calming hypothesis. Post-performance interviews with participants shed some light on the unpredicted behavior of the solo meditation. While the idea of facing a wall immediately on entering the space was meant to evoke privacy and to emulate the practice in some forms of Zen meditation, Zen meditation was not a strong, easily accessed reference point for many participants. Instead, the specific set up evoked in some a threatening situation with something unknown happening behind them. A few people also mentioned references to childhood punishment, as exemplified by “time out,” a widely used parenting practice in the last decades, adding to the stressful associations in this specific treatment for a minority of participants. Thus while post performance conversations revealed that attention focusing occurred during the solo meditation treatment, the effect was not as calming for some participants.

## Discussion

The analysis presented here supports the thesis that calming practices will have the effect of reducing parochialism and increasing altruism. Group meditation was found to predict higher levels of cooperation generally and to have an even larger effect of reducing parochialism. While the solo meditation treatment was not found to increase altruism, it was found to decrease parochialism. While Kang et al (2014) and demonstrate that LKM leads to decreases in parochialism in terms of racial bias, it is measured through reported attitude. Likewise, Lueke and Gibson (2014) demonstrate reduced age and race bias in association with increased mindfulness through implicit bias. This study measure of parochialism more actively through monetized decisions in PG games. Also LKM, as used in Kang et al, confounds relaxation effects with priming for universal compassion. This study demonstrates that such a reduction in parochialism happens independent of such priming. It has also been noted that experiments with mindful breathing practices potentially confound calming with attention focusing, and experiments suggest that attention focusing has effects on emotion regulation beyond calming (Arch & Craske, 2006). However, the treatment here less strongly emphasized attention re-focusing as part of the meditation than mindful breathing exercises in other experiments and added emphasis on whole body relaxation to reduce such confound. This study also uniquely looks simultaneously at general altruism and parochialism in order to clearly differentiate these effects. The use of a community sample that expanded on typical psychology experiments on undergraduates is another strength.

It is unclear from the experiment whether the difference in effect from the solo vs group variants had to do with the effects of doing the practice in private or if it had to do with inadvertently different effects on calm produced by idiosyncrasies of the experimental set up. The unsolicited reports from some participants about stressful associations with facing a wall indicate the usefulness of repeating the experiment with either a more uniformly comfortable way of creating privacy for the participants or an independent measure of calm after treatment.

As the winning in the game involved a random draw and the player’s choice affected the probability of winning, a potential challenge might be that an alternative interpretation of the impacts on altruism is that the effects might be mediated by a decrease in risk aversion, rather than other-regard. It is still, however, a relative evaluation of the risk of oneself winning vs probability of others winning and so game play should still be interpreted as a measure of altruism. Also, this critique would not extend to findings with regards parochialism, given that this is measured via differences between in and out-group play.

While there is a great deal of experimental and quasi experimental support for theories of ritual that predict that ritual practices performed together bond the group and lead to parochial altruism (Norenzayan & Shariff, 2008), this and other studies of meditation indicate that this is not true of all ritual practices performed in groups. It would be naïve to assume that the heterogeneous physical practices that have been traditionally lumped together into the category of ‘ritual’ should have identical effects, so this should not be a surprise. We should not be aiming for a single theory of rituals, but instead should use this loose category of activities that have been called ritual as a grab bag in which to find specific phenomena to study: synchronizing activities, activities which involve sacrifice, activities which involve calming, activities which involve contemplation of morally concerned high gods, etc. (Frost, 2013).

Durkheim’s definition of ‘ritual’ refers to their causal opacity (Durkheim, 1912). It is the category of social phenomena anthropologists use for activities whose material function is non-obvious to the anthropologist, an outsider. As Rappaport noted however, the Papua New Guinea tribesman who initiates a dance before a raid has a clear idea of the function of the ritual to generate group commitment (Rappaport, 1999). Similarly, centuries of Buddhists have sat in meditation in part because they observe that it cultivates compassion. As Rappaport writes, many of these disparate things which anthropologists have chosen to call ‘ritual’ are not causally opaque to their practitioners, whose beliefs about their function are actually accurate. The behavioral experiments we do now as scientists are in some cases likely replicating experiments done and quasi-experiments observed by people thousands of years ago.

Of course, a behavioral tendency toward altruism with non-group members presents an evolutionary conundrum. If altruists freely associate with others, they will be subject to problems with free-riders: people who take advantage of the altruist and fail to reciprocate. This suggests that such a behavioral tendency could be associated with some other compensating mechanisms. These could include independent physical or mental health benefits, positive assortment with fellow altruists, compensating horizontal cultural transmission (learning from unrelated others), such as through increased effectiveness in proselytizing and conversion. In different cases all three of these mechanisms have been shown to be at play for meditation. The empirical support for mental and physical health benefits of meditation, yoga, and related practices is now copious (Grossman, Niemann, Schmidt, & Walach, 2004) (Sloan, Bagiella, & Powell, 1999)(Koenig & Larson, 2001). While many religions profess universal altruism and adherents may ‘practice what they preach,’ in many of these cases, the groups practice high degrees of positive assortment. They may have an inclination to cooperate altruistically in all interactions, but they interact preferentially with co-religionists, who ‘coincidentally’ practice the same altruism-cultivating rituals (Wilson, 2002). For example, Christians during the Roman empire who actively practiced universal compassion tended to associate preferentially with each other for spiritual reasons and had special rules for the treatment of “brethren in poor standing” (non-cooperators), which escalated to exclusion. This represents a potential question for future experiments: “do calming practices cause people to preferentially associate with other calm people?” Wilson also reviews documentation of significant early Christian charity toward less fortunate non-Christians. This altruism was often associated, however, with conversion activity, whether it be active proselytizing or others joining the fold motivated by the greater material success and health of those in the fold. A Buddhist parallel would be association into sangha. While these circles of spiritual fellowship may be motivated by intellectual and spiritual exchange, it is plausible that they will also have material repercussions in terms of increased opportunities for economic coordination and mutual altruism. Of course, such evolutionary arguments assume evolutionary equilibrium, and given the dynamic nature of human culture, there is no reason to assume that such assumptions should be accurate. For this reason, such evolutionary arguments should simply be used to suggest research questions and should not be taken as claims about the state of the world; empirical determination of the actual effects of a behavior should be considered a separate question.

A strength of isolating the element of calming allows for an understanding of its independent effects. However, the limitation is that this then does not get at potential synergistic effects. In terms of the effects on social structure and therefore on dynamics of cultural group selection, there are parallels to be drawn to Harvey Whitehouse’s multiple modes theory of ritual (Whitehouse, 2002). If meditation and similar calming activities do indeed facilitate a more generalized altruism, they would also facilitate cooperation in larger groups. It could be that these doctrinal mode practices combine calming activities synergistically with other shared actions facilitating bonding with abstract social identifiers to bind large polities into cooperative units. In noting the contradictory evidence of intentional synchrony as a facilitator of parochialism, it may be that there is some synergy with arousal, dysphoria. Perhaps an explanation for the cases where such synchrony led to general prosociality is that the synergy with calm (post exercise high) as opposed to dysphoric arousal eliminated the parochialism and left an altruistic response. The question of such synergies amongst calming, synchrony, and/or arousal and dysphoria remains open. The importance of looking at such synergies has been noted already n the experimental literature on the effects of meditation (Holzel et al., 2011). For example, calm and non-reaction may synergize with explicit direction to reevaluate emotional responses by giving the time to enable such, and this could then lead to prosocial responses that are greater than would be anticipated from our understanding of their effects in isolation. Also looking at the anthropological literature, Rappaport notes the prevalence of reconciliation rituals amongst many societies when conflict needs to be resolved. Calm activities like sharing food are a frequent part of such practices and it may be that parochialism reducing effects of calming may be an essential ingredient in these practices. Again, this is an open question for experimentation.

There are obvious limits to this kind of experiment by itself. The treatments, of similar duration as other one-off treatments in experiments, were short and the nature of controlled experiment is that gains in precision are often at the expense of ecological validity: the ability to extrapolate those results to daily life. However, in synthesis with other experimental and survey based work on meditation and related calming practices, there is growing support developing for the theory that calming practices benefit prosociality, increasing altruism and reducing existing parochialism. Of course another limitation is that this study examines mean effects and does not look at the potential structure of variance of response in the population.

A few directions of further research are suggested. First, it would be useful to see if similar effects of reducing parochialism could be seen in standing identity groups rather than in the less stable temporary groups of an experimental set up with random group assignments. While this study demonstrates a reduction in parochialism due to calming practices, a reduction in parochial preference across standing identity lines of ethnicity, religion, or nationality would be evidence of a stronger, more ecologically valid, effect. Second, to address the evolutionary dilemma of altruism, it would be useful to examine the ethnographic record for correlations between calming ritual practices and positive assortment or increased conversion rates which maintain these rituals in the population in the face of potential free riders. There is already evidence of health benefits which may be sufficient to compensate for such potential losses, but there may be multiple factors contributing to the stability of such practices in the population in the face of free rider problems. Third, it would be useful to compare different ritual forms, including ones iconic of costly signaling or synchrony theories, in the same experimental arrangement as calming practices to test different theories of ritual social function. Fourth, it would be useful to tease apart the effects of different elements of meditation practice. Does attention focusing have effects on prosociality independent of calming? This could explain the difference between the effects of solo vs group meditations in this experiment, given that they both successfully focused attention but varied in their calming effects, as evidenced in post-experiment feedback. Further, does meditation on concepts like compassion, emptiness, or reduction of suffering have independent effects from physical calming and/or interaction effects and are the prosocial effects of calming practices direct or mediated by increased accuracy in social cognition, increase in compassion, or increased self regulation of emotional responses. Answering these questions will help us develop more sophisticated sciences of ritual practice and meditation.

### Ethical Considerations

All procedures performed in studies involving human participants were in accordance with the ethical standards of the institutional and/or national research committee and with the 1964 Helsinki declaration and its later amendments or comparable ethical standards. This experiment was carried out with the review and prior approval of the UC Davis Institutional Review Board. Informed consent was obtained from all individual participants included in the study.

No conflicts of interest exist.

## Acknowledgements

**Funding:** This study was conducted with funds provided through discretionary funds from the Department of Anthropology at UC Davis and with space provided by the Department of Theater and Dance at UC Davis as part of support for the authors MFA thesis work.

## Supplementary Material

### Treatment Group Scripts

Treatment Group A *(Control Group, no preparation)*
“You are going to be asked to play the Game directly. Please grab an instruction set, which we will read together.”

Treatment Group B *(Control Group, no directed preparation, but sharing a physical space for 5 minutes)*
“Welcome. Please feel free to wander the space for a bit, make yourself comfortable. You do not have any specific instructions. We will come together in a few minutes to take a survey.”
*(Instructions for facilitator: 5 minute pause.)*
“We are now going to play a game. Please come get an instruction set, which we will read together.”

Treatment Group C *(“Individual Ritual”)*
“Please make yourself comfortable somewhere in the space standing and facing one of the walls with plenty of personal space around you. Make yourself comfortable, with knees slightly bent, breathe easily.
We invite you to close your eyes and relax.
Feel your body and your breath.
Allow tension to drop away from your shoulders and any other place where it is not needed to keep you standing. Allow yourself to be comfortable. If you need to adjust yourself somehow, feel free.
Keep bringing your attention back to your breath. And allow tension to drop away with each inhale and exhale.
Please remain for 3 minutes here in this quiet awareness of your breath, feeling your body.
Again, adjust your stance as needed to make yourself comfortable.”
*(Instruction to facilitator: take 3 minutes in silence)*
“Keeping this sense of relaxation, please open your eyes. As you are ready, please come and get an instruction set for the Game, which we will read together.”

Treatment Group D *(“Group Ritual”)*
“Please come into a loose circle, standing in the center of the space. Make yourself comfortable, with knees slightly bent, breathe easily.
Take a look around for a few moments to see each other.”
*(Instruction to facilitator: pause for 30 seconds or so for people to see each other, but not so long that it becomes awkward)*
“We invite you to close your eyes and relax.
Feel your body and your breath.
Allow tension to drop away from your shoulders and any other place where it is not needed to keep you standing. Allow yourself to be comfortable. If you need to adjust yourself somehow, feel free.
Keep bringing your attention back to your breath. And allow tension to drop away with each inhale and exhale.
Please remain for 3 minutes here in this quiet awareness of your breath, feeling your body.
Again, adjust your stance as needed to make yourself comfortable.”
*(Instruction to facilitator: take 3 minutes in silence)*
“Keeping this sense of relaxation, please open your eyes. As you are ready, please come and get an instruction set for the Game, which we will read together.”

### The Game Instructions

*(each participant has a copy to read. All read together as the instructions are recited by the facilitator. At the appropriate moment, after the instructions are read, the facilitator asks if there are any clarifying questions about how the game works and answers them.)*

This Game is a variation on one that is used regularly in economic and political science experiments. Your choices in the game are used to evaluate theories of human behavior.

In the Game, all players are given 10 points and a choice to keep some number of these points and place some number of these points in a Common Pool. There will be a cash prize won by one player in each round of the Game.

In each round, the more points contributed by all to the Common Pool, the larger the prize, up to $20. To be specific, the Prize goes up linearly with the number of points in the Common Pool. If everyone puts in 10 points, the prize is $20. If no one puts in points, there is no prize.

The more points that you keep for yourself, the greater your chance of winning the Pool. By analogy to a raffle, you have one raffle ticket per point that you keep. If you keep NO Points, then you have no raffle tickets. If everyone keeps the same number of points, everyone has the same chance of winning. If you contribute more points to the Common Pool, you decrease your chance of winning. If you put in less points, you increase your chance of winning.

You may also default to No Play. In this case, you contribute nothing to the Common Pool and have no chance of winning.

To repeat, the important things to remember are:

- the more points you contribute, the larger the prize.
- The more points you keep relative to others, the higher the likelihood of you winning the prize

*Your choice and other’s choices will be anonymous*.

‐‐‐‐

You will play the Game twice.

- In the first round, you will play with the other people in your Preparation Group.
- In the second round, you will play with a mixed group from other Preparation Groups, whom you will not meet.

Each of the two round starts again with a new 10 points and a new prize.

We do not specify a goal of the Game. There are just choices presented to you and outcomes that will result.

We would like you to understand the game before playing. If you do not understand how the game works, please ask one of the performers, and they would be happy to take a few moments to explain.

**Please do not discuss the game with each other during the experiment. Afterward, we encourage discussion! We would like to ask that you not discuss the game in detail with anyone whom you know to be attending the show in the future, until after they have participated.**

**The Game -**

**Your Answers (Group X), Player Number ________**

In each Round, you are given a new 10 points to play with and there is a new prize. The more points contributed by the group to the Common Pool, the larger the prize, up to $20. The more points you keep relative to others, the larger your chance of winning the prize.

**Round 1 You are playing with a group of players, all of whom are from Group X. Of your 10 Points, how many Points do you place in the Common Pool?**

(circle a number. The rest, you keep.)

**1 2 3 4 5 6 7 8 9 10 *No Play***

**Round 2 You are playing with a mixed group of players from all Groups. Of your 10 Points, how many Points do you place in the Common Pool?**

(circle a number. The rest, you keep.)

**1 2 3 4 5 6 7 8 9 10 *No Play***

‐‐

**At the end of the performance, the results will be given and briefly discussed, and the cash prizes will be given out.**

**Keep your number to claim your money if you win.**

**Demographic Information**

**This is a short survey to get some very basic demographic information. All information is anonymous and will just be used in analyzing variations in playing choices.**

**Age____________**

**Sex____________**

**Education Level (circle one):** High School, Some College, Bachelors Degree, Graduate Level Studies

**Relationship Status___________**

**Children___________**

**Have you participated in the performance before or the Community Workshop? (circle one) Yes No**

**Have you studied this kind of game structure before? Public Goods Game, Dictator Game, Ultimatum Game, etc? (circle one) Yes No**

**Any comments?**

